# Clonal evolution and specificity of the human T follicular helper cell response to *Plasmodium falciparum* circumsporozoite protein

**DOI:** 10.1101/2021.09.10.459751

**Authors:** Ilka Wahl, Anna Obraztsova, Julia Puchan, Rebecca Hundsdorfer, Sumana Chakravarty, B. Kim Lee Sim, Stephen L. Hoffman, Peter G. Kremsner, Benjamin Mordmüller, Hedda Wardemann

**Author notes:** Correspondence: Hedda Wardemann, German Cancer Research Center (DKFZ), Im Neuenheimer Feld 280, 69120 Heidelberg, Germany.

## Abstract

T follicular helper (T_FH_) cells play a crucial role in the development of long-lived, quality-improved B cell responses after infection and vaccination. However, little is known about their clonal evolution. Here we assessed the cell phenotype, clonal dynamics, and TCR specificity of human circulating T_FH_ (cT_FH_) cells at monoclonal level during successive malaria immunizations with radiation-attenuated *Plasmodium falciparum* (*Pf*) sporozoites. Repeated parasite exposures induced a dynamic, polyclonal cT_FH_ response with high frequency of cells specific to the *Pf* circumsporozoite protein (PfCSP), the main surface protein of sporozoites and a validated vaccine target. Repeated immunizations were required to induce detectable PfCSP-reactive cT_FH_ cell responses to a small number of epitopes. HLA-restrictions and differences in TCR generation probability explain the high targeting frequency of the polymorphic Th2R/T* region over the conserved T1 epitope. The vast majority of anti-Th2R/T* TCRs failed to tolerate natural polymorphisms in their target peptide sequence suggesting that parasite diversity limits natural boosting of the cT_FH_ cell response in endemic areas and protection from non-vaccine strains. Among convergent anti-Th2R/T* TCRs with high sequence similarity, subtle differences in CDR3 composition discriminated cross-reactive from non-cross-reactive cT_FH_ cells. Thus, our study provides deep molecular and cellular insights into the kinetics, fine specificity and HLA-restrictions of the anti-cT_FH_ cell response that are of direct relevance for the design of PfCSP-based malaria vaccines by guiding the selection of PfCSP peptides that induce optimal B cell help.

## INTRODUCTION

T follicular helper (T_FH_) cells play a key role in B cell selection and antibody affinity maturation during germinal center (GC) reactions and are hence essential for the generation of protective B cell responses^1,2^. Therefore, the targeted induction of strong T_FH_ cell responses is a major goal in the design of vaccines that aim at inducing protective humoral immunity. Clonal relatives of human GC T_FH_ cells are detectable in peripheral blood^3–5^. These circulating T_FH_ (cT_FH_) cells show strong transcriptional and phenotypic similarities with their GC counterparts including expression of the GC-entry chemokine receptor CXCR5^6,7^. Based on the expression of the chemokine receptors CXCR3 and CCR6 three subsets can be distinguished: cT_FH_1 cells (CXCR3^+^CCR6^−^) with lower helper cell capacity and cT_FH_2 cells (CXCR3^−^ CCR6^−^) and cT_FH_17 cells (CXCR3^−^CCR6^+^) with higher helper capacity^6,7^. A small fraction of cT_FH_ cells expresses activation markers like PD-1 and ICOS and closely resembles GC T_FH_ cells^3,4^. Thus, activated cT_FH_ cells can be used to assess the clonal evolution, cell phenotype, and epitope specificity of GC T_FH_ cells to optimize vaccine design strategies.

*Plasmodium falciparum* (*Pf*) malaria is a vector-borne parasitic disease for which a highly efficacious vaccine that induces durable protective immune responses is not yet available. Numerous vaccine candidates against *Pf* have been developed that target different stages of the complex life cycle. The most advanced *Pf* malaria vaccine, RTS,S AS01, targets *Pf* circumsporozoite protein (PfCSP), the major surface protein on sporozoites, the parasite stage that is transmitted to humans by the bite of infected *Anopheles* mosquitoes. Immunization with RTS,S can block the infection before symptom onset and has the potential to confer sterile protection^8^. However, efficacy is limited and protection only short-lived^9,10^.

To design a PfCSP-based vaccine with improved efficacy, the humoral immune response to PfCSP has been characterized in depth. Recent studies at monoclonal level shed light on the gene features and clonal evolution of protective anti-PfCSP antibodies and identified potent target epitopes that are not contained in RTS,S^11–14^. Similar to anti-PfCSP antibody titers, PfCSP-specific cT_FH_ cell responses have been shown to correlate with protection after RTS,S immunization^15^. However, studies characterizing the human PfCSP-specific cT_FH_ cell response at monoclonal level have not been performed. Thus, how PfCSP-specific cT_FH_ cell responses evolve and the molecular features and epitope specificity of their TCRs are not known.

Here, using cells from subjects immunized with aseptic, purified, cryopreserved *Pf* sporozoites (PfSPZ Vaccine^16–18^), we carried out functional TCR repertoire studies at single-cell level to define the clonal dynamics, epitope specificity, and MHC linkage profile of human PfCSP-reactive cT_FH_ cells by expression cloning of TCRs from activated cT_FH_ cells isolated at different time points after successive sporozoite immunization. The data provide a deep molecular and cellular understanding of the human anti-PfCSP T_FH_ cell response.

## RESULTS

### Activation and clonal expansion of circulating T_FH_ cells after successive PfSPZ immunization

To characterize the cT_FH_ cell response to *Pf*, we analyzed longitudinal peripheral blood samples from participants of a vaccine trial with radiation-attenuated NF54 sporozoites (Fig. 1a). Five malaria-naive individuals (D1 - D5) were immunized three times on days 1, 8 and 29 by direct venous inoculation of 9×10^5^ radiation-attenuated, aseptic, purified, cryopreserved *Pf* sporozoites (Sanaria PfSPZ Vaccine), which induced a robust antibody response against PfCSP (Fig. 1b). Peripheral blood mononuclear cells (PBMCs) were isolated three days before the first (0) and seven days after each of the three immunizations (I-III) with the exception of donor D5 for which sample (I) was not available. Upon immunization, the frequency of activated CD45RA^−^CD4^+^CXCR5^+^ cT_FH_ cells with PD-1 and ICOS expression increased strongly suggesting that these cells actively participated in the anti-sporozoite response (Fig. 1c-e). The majority of PD-1^+^ICOS^+^ cT_FH_ cells showed a CXCR3^+^CCR6^−^ cT_FH_1 cell phenotype compared to a minor fraction of CXCR3^−^CCR6^−^ cT_FH_2 cells, the dominant subset at study onset, and a small number of CXCR3^−^CCR6^+^ cT_FH_17 cells (Fig. 1f,g). An increase in the frequency of CXCR3^+^CCR6^−^ cells was also observed among activated CD4^+^ central memory (T_CM_) and effector memory (T_EM_) T cells (Supplementary Fig. 1). Thus, the overall anti-sporozoite CD4^+^ T cell response was dominated by cells with a T_H_1 phenotype.

**Figure 1:**
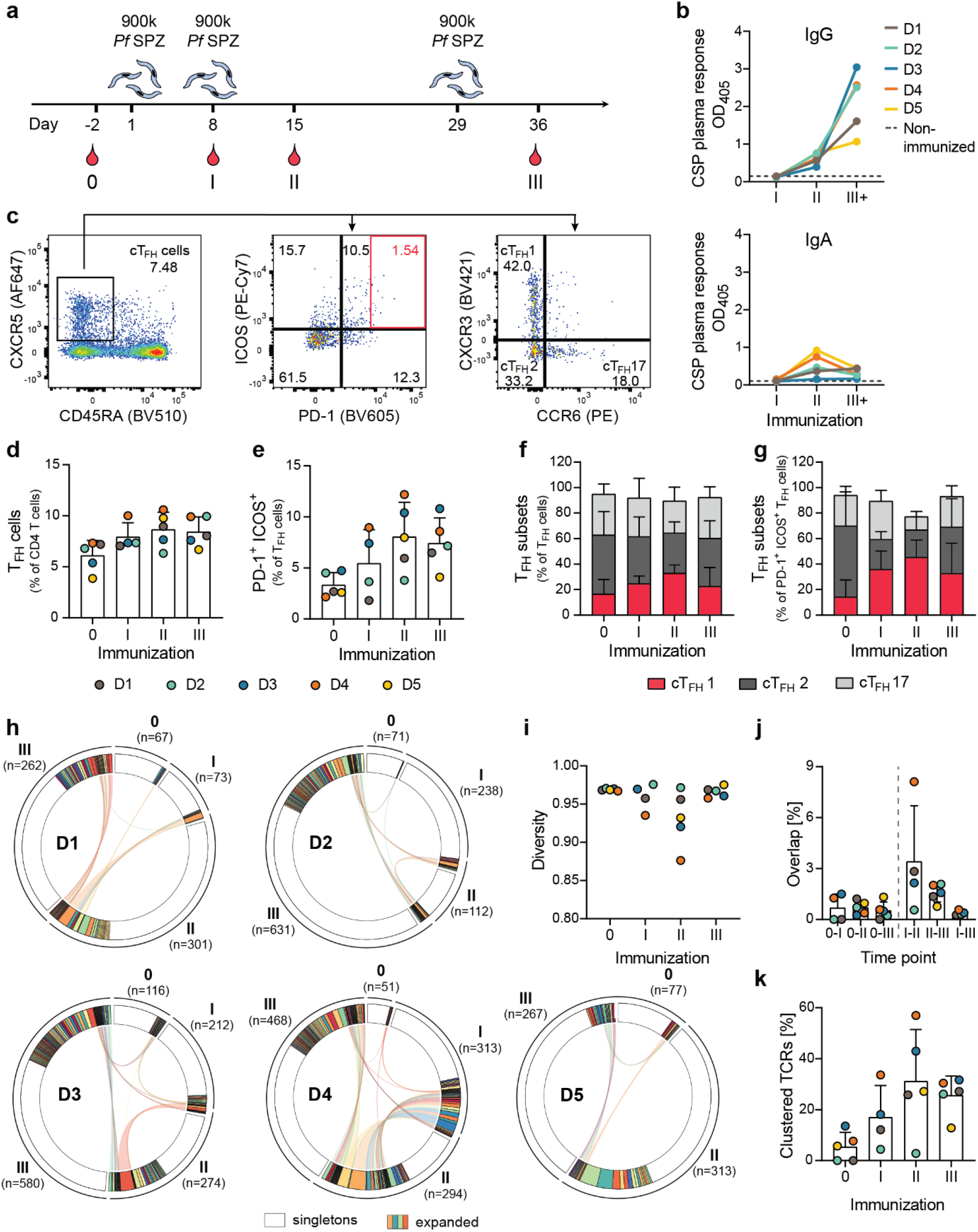
Induction of cT_FH_ cell activation and clonal expansion after PfSPZ Vaccine immunization. (**a**) PfSPZ Vaccine immunization scheme. Volunteers were immunized with radiation-attenuated PfSPZ at days 1, 8 and 29 and blood was sampled 3 days before the first immunization (0) and 7 days after each immunization (I, II, III). (**b**) IgG and IgA plasma antibody titers against PfCSP determined by ELISA. Colored lines represent individual donors, dashed lines indicate the mean OD_405_ of three non-immunized donors. One experiment representative of three independent experiments is shown. (**c**) Representative gating strategy for the flow cytometric analysis of PBMC samples, pre-gated on live CD3^+^CD4^+^ T cells (Supplementary Fig. 1a). The red square indicates the sorting gate of highly activated cT_FH_ cells for TCR repertoire analysis. **(d**,**e)** Percentage of cT_FH_ cells (CD3^+^CD4^+^CD45RA^−^CXCR5^+^) (**d**) and activated cT_FH_ cells (PD-1^+^ICOS^+^) (**e**). Bar graphs show mean and SD, dots mark individual donors. (**f**,**g**) Percentage of cT_FH_1 (CXCR3^+^CCR6^−^), cT_FH_2 (CXCR3^−^CCR6^−^) and cT_FH_17 cells (CXCR3^−^CCR6^+^) among all cT_FH_ cells (**f**) or activated cT_FH_ cells (**g**). The mean and SD are shown. **(h-k)** TCR sequence analysis of sorted PD-1^high^ ICOS^+^ cT_FH_ cells. Clones were defined by identical TRB V and J segment usage and identical CDR3 nucleotide sequence. (**h**) Circos plots showing clone dynamics over the course of immunizations. Expanded clones are shown in different colors. Clones overlapping between time points are connected with lines. Non-expanded clones are shown in white. *n* indicates number of TRB gene sequences. (**i**) Repertoire diversity quantified by normalized Shannon-Wiener index. (**j**) Repertoire overlap between time points quantified as a fraction of shared clones. (**k**) Percentage of convergent TCRs as determined by TCR clustering (Supplementary Fig. 3 and 4). Bar graphs show mean and SD, dots mark individual donors (**j**,**k**). (d-k) A sample from time point I was not available.

To enable the unbiased integration of cell phenotype data with TCR gene information, we used indexed flow cytometric cell sorting to isolate single PD-1^high^ICOS^+^ cT_FH_ cells from all sampling time points and amplified and sequenced the paired TRB and TRA genes from over 3,000 cells without antigen-mediated enrichment or prior antigenic stimulation^19^. Upon immunization, cT_FH_ cells expanded clonally resulting in a drop in repertoire diversity quantified by normalized Shannon Wiener index (Fig. 1h,i). The degree of expansion was highest after the second immunization, presumably because of the shorter time interval of only seven days between the first and second compared to 21 days between the second and third dose. Samples from neighboring time points tended to have a higher clonal overlap, although only a few expanded clones persisted over multiple time points indicating that the cT_FH_ cell repertoire composition changed with time (Fig. 1j).

Although the TCR gene usage was highly heterogeneous (likely reflecting the antigenic diversity of the parasite with over 5,000 genes^20^), we identified several TRBV and TRAV genes in individual donors whose usage frequency increased over time indicating that cT_FH_ cells expressing these segments contributed to the anti-sporozoite immune response (Supplementary Fig. 2). Furthermore, numerous TCRs with high sequence similarity were detected in different donors suggesting that cT_FH_ cells expressing these convergent TCRs, which increased in frequency upon immunization, recognized the same sporozoite antigen (Fig. 1k).

In summary, successive PfSPZ Vaccine immunizations induced a polyclonal and highly dynamic cT_FH_ cell response characterized by signs of antigen-mediated TCR selection.

### PfCSP-specific cT_FH_ cells emerge late and predominantly target the Th2R/T* region

To determine whether cells that showed signs of *Pf*-driven selection (clonal expansion, expression of V segments with increased usage frequency upon immunization or TCR convergence) included PfCSP-reactive cT_FH_ cells, we selected a representative set of 245 T cell clones for functional TCR analyses (Supplementary Table 1). Through gene cloning of paired TRA and TRB genes and subsequent retroviral transduction, we stably expressed the selected TCRs in CD3^+^CD4^+^, endogenous TCR-negative Jurkat76 cells (Supplementary Fig. 5)^19^. To screen for PfCSP reactivity, the cell lines were co-cultured with EBV-immortalized autologous B cells pulsed with pools of overlapping 15mer peptides covering the complete N-terminus and NANP repeat region or the C-terminus (hereafter referred to as N-CSP or C-CSP, respectively; Supplementary Table 2). Using IL-2 secretion as read-out for T cell activation, we identified 57 N- or C-CSP specific clones demonstrating that nearly one fourth of all tested cT_FH_ cell TCRs (23%; 57/245) recognized PfCSP (Fig. 2a,b). The majority responded to C-CSP peptides (82%; 47/57) and were identified in all donors at a frequency of 9-34%. In contrast, the relatively small number of N-CSP reactive TCRs (18%; 10/57) were obtained from only one donor (D3). The data highlight the strong immune activating capacity of PfCSP compared to other sporozoite antigens and relative immunodominance of the C-terminal compared to the N-terminal domain.

**Figure 2:**
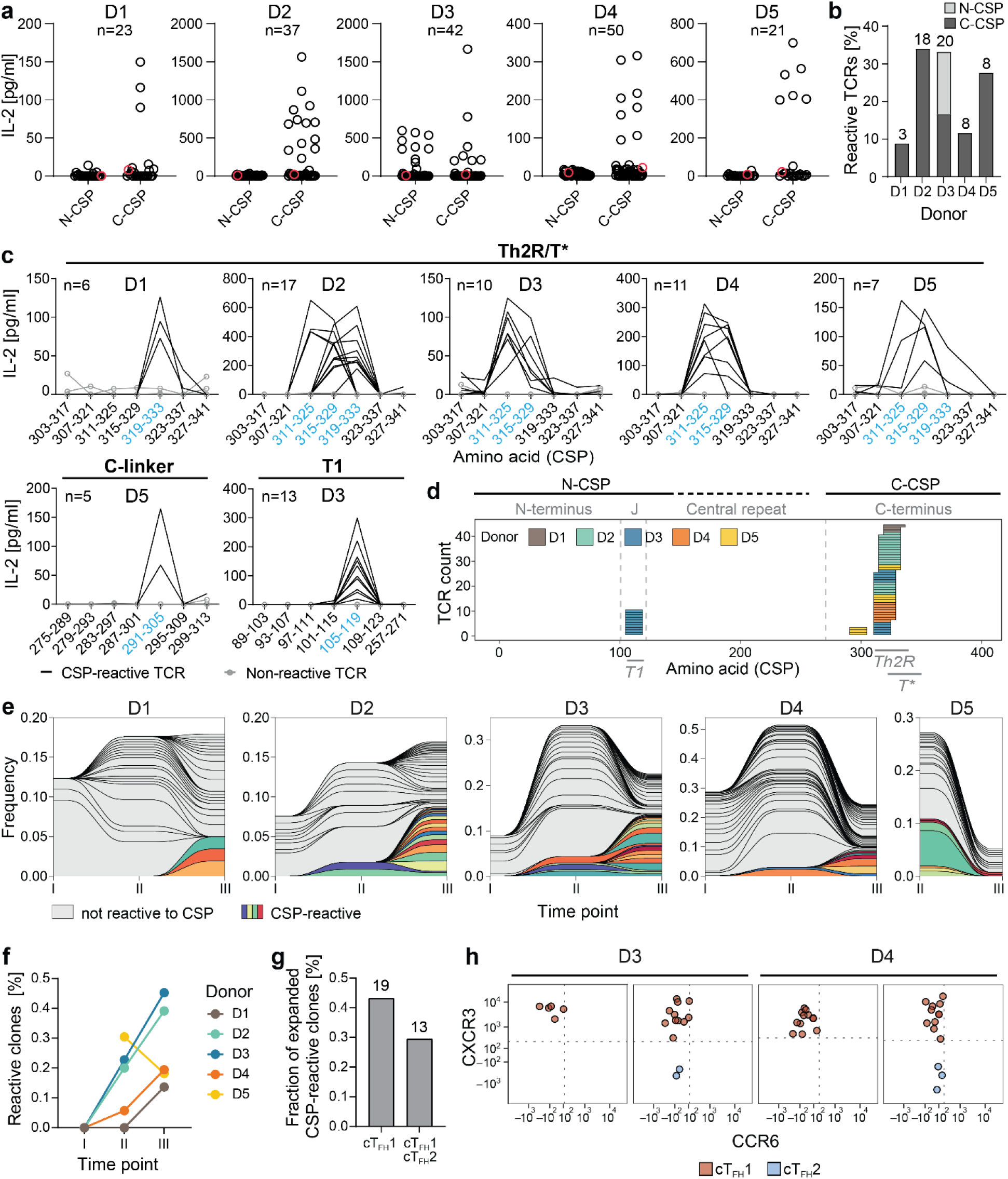
PfCSP-reactive TCRs are strongly biased to the Th2R/T* region and emerge late. (**a-d**) TCR-transgenic Jurkat76 T cells were co-cultured with autologous B cells pulsed with N-CSP or C-CSP peptide pools (**a**,**b**) or single peptides (**c**,**d**). T cell activation was quantified by IL-2 ELISA. (**a**) IL-2 values for a selected set of TCRs, each dot shows an individual TCR, the negative control TCR is highlighted in red. (**b**) Frequency of PfCSP-reactive TCRs among tested. The total number of reactive TCRs per donor is indicated. (**c**) Selected set of TCR-transgenic T cells stimulated with single peptides covering different PfCSP epitopes. Each line shows an individual TCR. The negative control TCRs (3) are shown in gray. The total number of tested TCRs is indicated. (**d**) Summary of all epitopes targeted by PfCSP-reactive TCRs. Regions of PfCSP covered by the N-CSP or C-CSP peptide pools are indicated. The central repeat region is covered by a limited number of peptides capturing all possible repeating motif combinations (dashed line). Published epitopes are indicated in gray below. (**e**) Kinetics of clonally expanded cells in the cT_FH_ cell repertoire of each donor. Only cT_FH_ cell clones of tested TCRs are shown. PfCSP-specific clones are highlighted in colors. Fraction of PfCSP-reactive T cell clones among tested. (**g**,**h**) Integration of flow cytometric index data. Frequency of expanded PfCSP-reactive clones with cT_FH_1 only or cT_FH_1 and cT_FH_2 phenotype. (**h**) Representative PfCSP-reactive clones with cT_FH_1 only or cT_FH_1 and cT_FH_2 (CXCR3^−^ CCR6^−^) phenotype. Each dot marks a single T cell. (**a-d**) show results from one representative out of three independent experiments. (e-f) A sample from time point I was not available.

To define the fine specificity of each PfCSP-reactive TCR, we performed stimulation assays with individual peptides. C-CSP reactivity was mostly limited to three overlapping peptides spanning amino acids (aa) 311-333 (PSDKHIKEYLNKIQNSLSTEWSP), overlapping with the known T cell epitopes Th2R and T* (Fig. 2c)^21–23^. In total, 44 TCRs from all donors showed reactivity to aa 311-333, hereafter referred to as Th2R/T*, and often responded to more than one of the overlapping peptides. In contrast, aa 291-305 (RNVDENANANSAVKN), represented by a single peptide in the C-terminal linker region, were recognized by only three TCRs from a single donor (D5). Similarly, all N-CSP reactive TCRs from D3 recognized a single peptide spanning aa 105-119 (PNANPNVDPNANPNV), a conserved region in the N-terminal junction containing a known CD4^+^ T cell epitope referred to as T1^24^. TCRs with reactivity to repeating NANP-motifs were not identified. Thus, the anti-PfCSP specific cT_FH_ cell response was limited to only three epitopes and targeted predominantly the Th2R/T* region (Fig. 2d).

Despite their relative abundance, PfCSP-reactive cT_FH_ cells emerged only after the second immunization, were rarely shared between sampling time points, and were most abundant after the third immunization (Fig. 2e,f). The fact that PfCSP-reactive cT_FH_ cells were not detected among the large expanded clones that were isolated after the first immunization suggests that these cells might recognize other *Pf* antigens or represent non-*Pf* specific bystander cells. The majority of PfCSP-reactive clones showed a cT_FH_1 phenotype (Fig. 2g,h). However, numerous clones were not phenotypically restricted and often included cT_FH_1 and cT_FH_2 cells suggesting that cells from the same clone exert different effector functions.

Thus, despite the high frequency of anti-PfCSP cT_FH_ cells after PfSPZ Vaccine immunization, the cells emerged relatively late in the response, showed a cT_FH_1-biased phenotype, and targeted a limited number of PfCSP epitopes with strong focus on aa 311-333 in the Th2R/T* region in most but not all individuals.

### MHC-II genotype limits the breadth of the PfCSP-specific cT_FH_ cell response

To determine whether the high frequency of anti-Th2R/T* TCRs and sparsity of anti-T1 and anti-C-linker TCRs were linked to presentation on different HLA molecules, we used anti-DP, -DQ or -DR antibodies to block TCR-pMHC binding specifically for each HLA locus (Fig. 3a). With two exceptions, Th2R/T*-specific TCRs in all donors recognized their target epitope in the context of HLA-DR, while all T1 or the C-linker specific TCRs recognized the peptides in the context of HLA-DQ (Fig. 3b). Gene sequencing-based HLA genotyping showed that nearly all donors shared specific HLA alleles with strongest overlap in the HLA-DR locus (Fig. 3c,d). We therefore investigated whether Th2R/T*-specific TCRs recognized their target peptides in the context of these shared HLA-DR alleles by stimulating the TCR-transgenic T cells with either autologous or heterologous peptide-pulsed B cells from each individual (Fig. 3e). These B cell swapping experiments showed that all Th2R/T*-specific TCRs from D3 could be stimulated by B cells from D4 and vice versa, but not by B cells from any of the other donors (Fig. 3f). Thus, the Th2R/T*-specific cT_FH_ cells from D3 and D4 likely recognize their target peptides in the context of the shared HLA molecules DRB1*15:01P or DRB5*01:01P (Fig. 3d,f; Supplementary Table 3). Similarly, all HLA-DR restricted Th2R/T*-specific TCRs from D2 and D5 responded to stimulation with pulsed B cells from both individuals suggesting that they bound their target peptides presented on the shared DRB1*04:02P HLA molecule. The two HLA-DR restricted Th2R/T*-specific TCRs from D1 reacted to peptide-pulsed B cells from D2 and D3, presumably in the context of the shared HLA-DR allele DRB1*07:01P. However, the opposite was not true. All TCRs from D2 or D3 were restricted to other HLA alleles reflecting a strong preference for non-DRB1*07:01P mediated HLA presentation of Th2R/T* peptides in these donors (DRB1*04:02P in D2 and DRB1*15:01P or DRB5*01:01P in D3).

**Figure 3:**
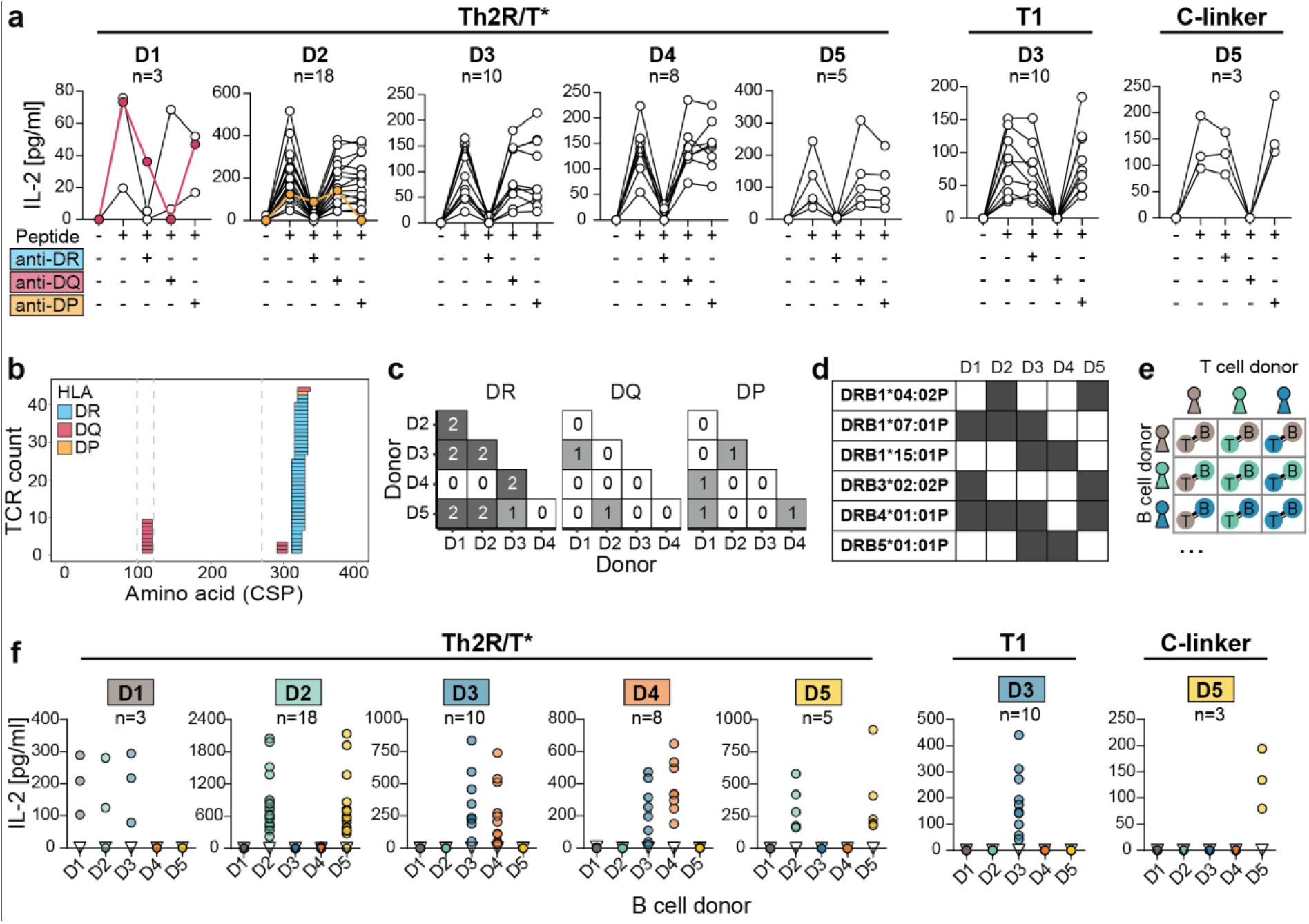
Th2R/T* (aa 311-333) immunodominance due to presentation on diverse HLA-DR alleles. (**a**) Stimulation of Th2R/T*, T1 or C-linker-specific TCR-transgenic T cell lines with autologous B cells in the presence (+) or absence (-) of anti-HLA-DR, -DQ or –DP blocking antibodies. Dots connected by lines indicate individual T cell lines. DQ- and DP-reactive anti-Th2R/T* TCRs are highlighted in red and orange, respectively. (**b**) Summary of HLA-blocking experiments. Colored rectangles represent the PfCSP peptide reactivity profile for individual DR (blue), DQ (red) and DP (orange) specific TCRs. (**c**) Number of shared MHC-II allele combinations across donors. (**d**) List of HLA-DR alleles shared between donors. Dark gray marks the presence of an allele in the respective donor. (**e**,**f**) B cell swapping experiment. (**e**) Graphical illustration of the experimental layout. (**f**) IL-2 secretion by TCR-transgenic T cell lines from the indicated donors after stimulation with autologous or heterologous B cells loaded with the indicated peptide. The number of tested TCRs is indicated. Triangles mark data from a non-PfCSP reactive control TCR. (**a**,**b**,**f**) One experiment representative for three independent experiments is shown.

Activation of the anti-T1 and C-linker TCRs from D3 and D5, respectively, was limited to autologous B cells, indicating that they recognized their target epitopes presented on non-shared HLA DQ alleles. To validate our experimental findings and to define the HLA alleles with higher likelihood of presenting the cT_FH_ cell target peptides in each donor, we used and compared four established computational prediction tools (NetMHCIIpan-4.0^25^, Sturniolo^26^, MixMHC2pred^27^ and MHCnuggets^28^; Supplementary Table 3). Unfortunately, the high discrepancy among the results and inconsistency with the experimental data made accurate predictions impossible.

In summary, the immunodominant aa 311-333 peptides in the Th2R/T* region are presented predominantly on shared HLA-DR alleles, while TCRs against the T1 and C-linker epitope recognize the peptide in the context of non-shared HLA-DQ alleles. The cT_FH_ cell response in individual donors shows a bias towards preferential recognition of specific pMHC complexes.

### TCR sequence features associated with PfCSP binding

To investigate whether certain TCR properties were associated with binding to different PfCSP epitopes, we analyzed the aa sequence features of all PfCSP-reactive TCRs. Th2R/T*- and T1-specific TCRs utilized a diverse set of TRAV and TRBV segments (Fig. 4a). Nevertheless, compared to non PfCSP-reactive TCRs, *TRBV20-1* was overrepresented among Th2R/T*-reactive TCRs in D2 and D5 and *TRBV30* was enriched among T1-specific TCRs from D3. (Fig 4b). Numerous *TRBV20-1* TCRs from D2 and D5 that recognize Th2R/T* presented on DRB1*04:02P MHC complexes belonged to convergent TCR clusters (Fig. 4c, Supplementary Table S4). Although clusters of convergent TCRs were only detected in donors in which the respective target peptides were presented on shared HLA complexes, *TRBV20-1* was also utilized by Th2R/T*-reactive cT_FH_ cells from D1 and D4 without TCR similarity, suggesting that *TRBV20-1* usage might play a role in the recognition of Th2R/T* peptides beyond DRB1*04:02P MHC complexes. In contrast to the similarity among *TRBV20-1* anti-Th2R/T* TCRs, T1-specific TCRs including those utilizing *TRBV30* were highly diverse and did not form clusters of convergence (Fig. 4c). To investigate the TCR repertoire limits beyond segment usage, we estimated the generation probability of PfCSP-specific TCRs. Strikingly, anti-Th2R/T* TCRs had a higher generation probability compared to the overall TCR repertoire and anti-T1 TCRs (Fig. 4d).

**Figure 4:**
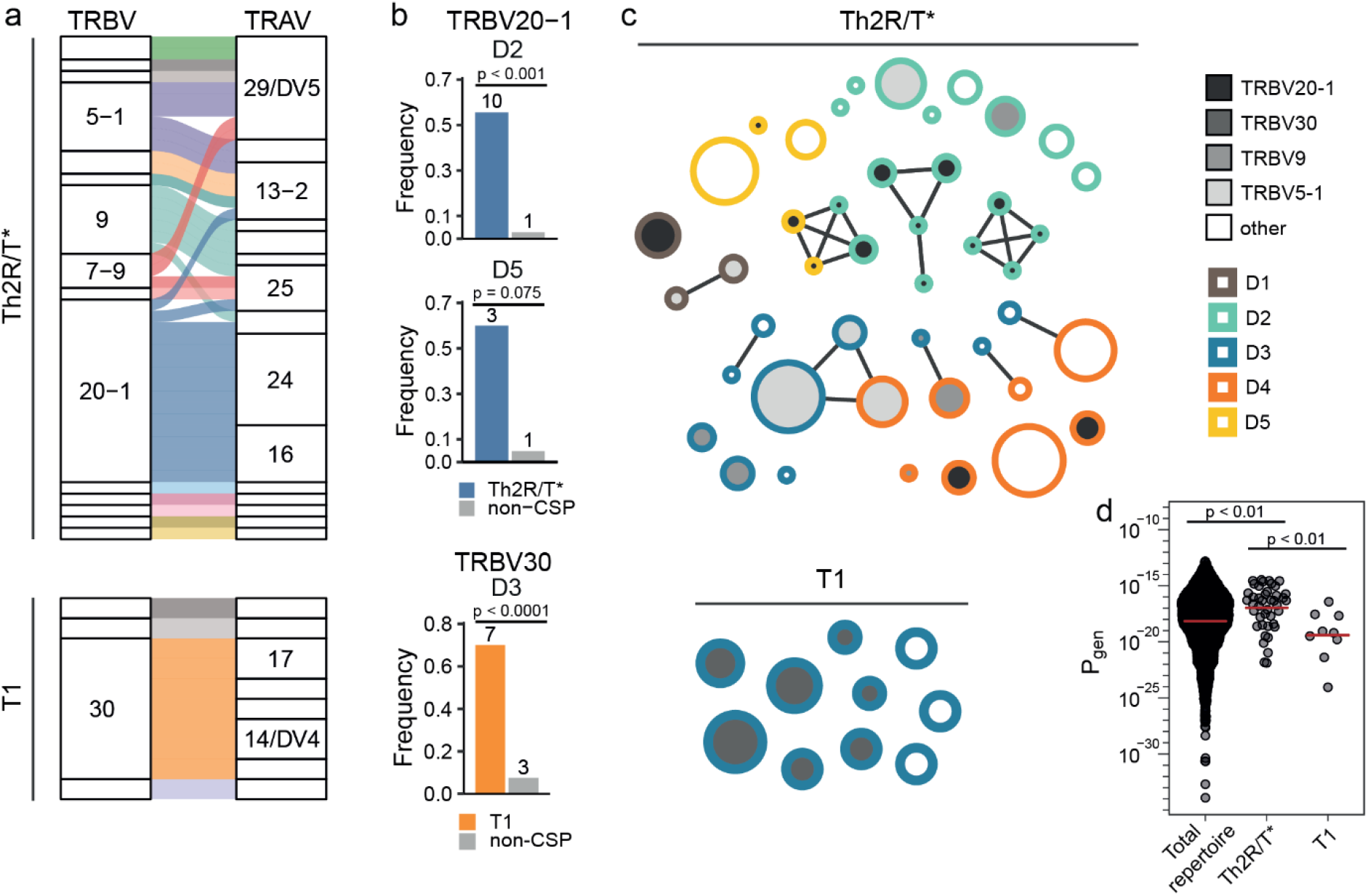
*TRBV20-1* and *TRBV30* are linked to Th2R/T* and T1-binding, respectively. (**a**) *TRBV* segment usage of TCRs recognizing the Th2R/T* (upper panel, n=44) and T1 (lower panel, n=10). (b) Frequency of *TRBV20-1* or *TRBV30* usage of Th2R/T* or T1-specific TCRs, respectively, compared to non PfCSP-specific TCRs in individual donors. (**c**) Clustering of TCRs based on sequence similarity. Each circle depicts an individual T cell clone and clones with convergent TCRs are connected by lines. Circle size represents clone size. (**d**) Generation probability of TCRs calculated using OLGA^51^. p values were calculated by chi-square test (**b**) and two-sided Mann-Whitney test (**d**).

Thus, despite the overall high TCR diversity of the anti-PfCSP cT_FH_ cell response, *TRBV20-1* and *TRBV30* seem to be associated with Th2R/T* and T1 specificity, respectively. The high generation probability of TCRs targeting aa 311-333 in the Th2R/T* region may contribute to the bias of the anti-PfCSP cT_FH_ cell response towards this region.

### Limited cross-reactivity of Th2R/T*-specific TCRs

Since the Th2R/T* region is highly polymorphic, we investigated how the degree of sequence diversity observed in natural *Pf* populations affects T cell activation by stimulating all TCR clones with autologous B cells that had been pulsed with a peptide covering aa 311-333 of the NF54 vaccine strain or of three representative *Pf* strains (7G8, K1, Dd2) from different geographic locations (Fig. 5a,b). None of the TCRs showed cross-reactivity to 7G8 or K1, which differ from NF54 by five and four aa mismatches, respectively, and only a limited number of TCRs (7/18) recognized the Dd2 peptide (Fig. 5c). All Dd2 cross-reactive TCRs were specific to aa 319-333 in the C-terminal part of the Th2R/T* epitope with only one aa mismatch compared to NF54 (Fig. 5b). Donors D3 and D4 without cT_FH_ cells that recognized aa CSP319-333 lacked cross-reactive TCRs (Fig. 5c,d). Differences in the targeting preferences of peptides in the Th2R/T* region were linked to HLA-DR differences and were similar in donors with shared alleles (D2 and D5, D3 and D4; Fig. 5d), suggesting that HLA genotype restrictions impact the ability to generate NF54-Dd2 cross-reactive TCRs. However, only 7/18 TCRs that responded to the NF54 PfCSP aa 319-333 peptide showed Dd2 cross-reactivity demonstrating that even a single aa mismatch abrogated binding to the pMHC complex for the majority of TCRs (Fig. 5e). Cross-reactivity was not linked to gene usage and even members of convergent TCR clusters with identical V and J segment usage in their TRA and TRB genes differed in their cross-reactivity (Fig. 5f,g). Thus, NF54-Dd2 cross-reactivity is limited to cT_FH_ cells with TCR reactivity to aa CSP319-333 in the Th2R/T* region. However, minor CDR3 aa differences define the TCR cross-reactivity capacity of cT_FH_ cells and their ability to recognize aa CSP319-333 peptide variants that differ by as little as one aa from the NF54 vaccine strain.

**Figure 5:**
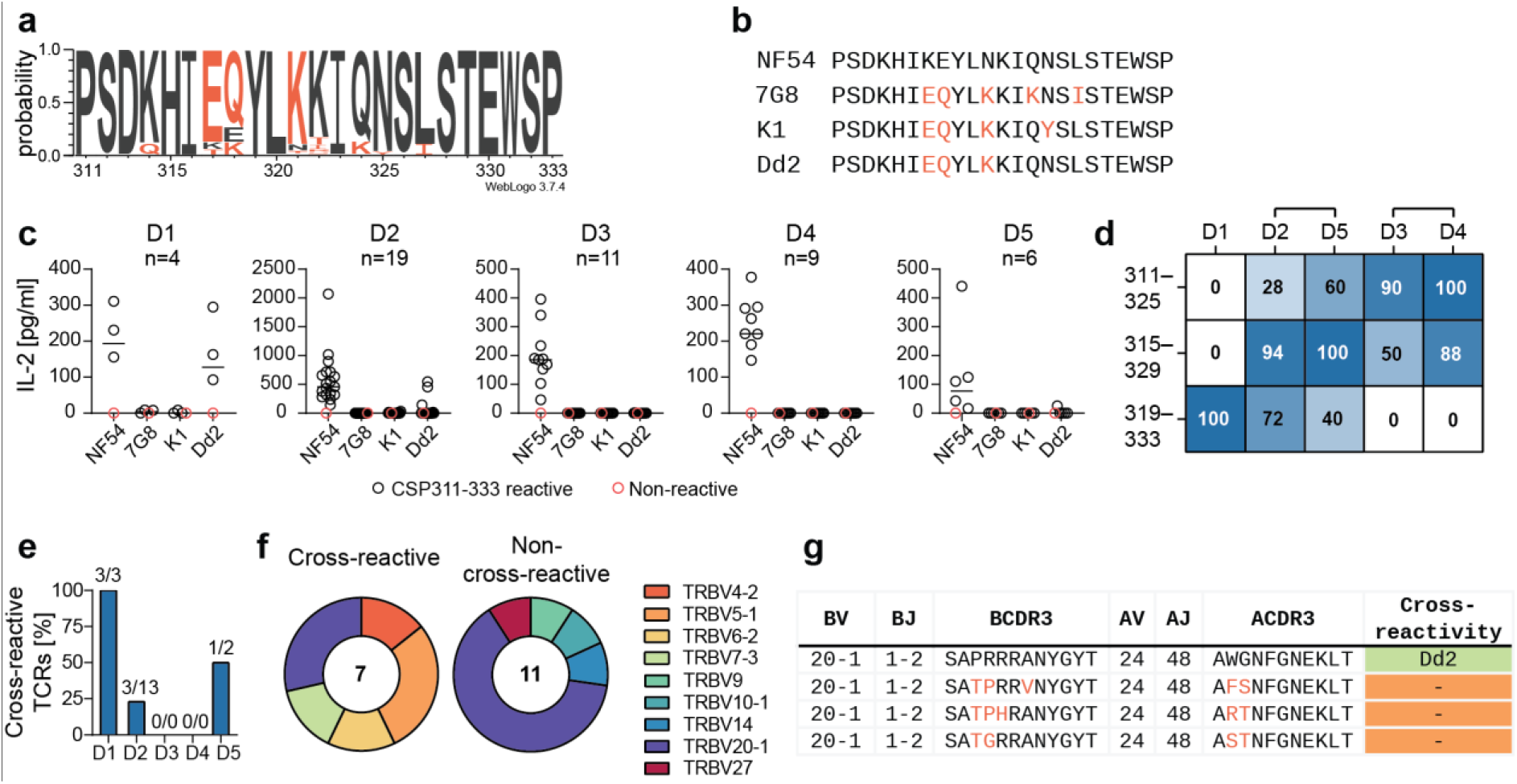
Limited cross-reactivity of Th2R/T* (aa 311-333) specific TCRs to common PfCSP variants. (**a**) Alignment of PfCSP aa 311-333 from 979 *Pf* isolates^58–60^. Amino acids corresponding to the sequence of the immunizing *Pf* strain NF54 are shown in dark gray, orange highlights amino acids differing from NF54. (**b**) Th2R/T* (aa 311-333) peptides from *Pf* strains NF54 (P19597), 7G8 (AB121015.1), K1 (AJ269946.1), Dd2 (AB121017.1). Mismatches compared to NF54 are highlighted in orange. (**c**) IL-2 secretion of TCR-transgenic T cell lines (black circles) after stimulation by autologous B cells pulsed with the indicated Th2R/T* (aa 311-333) peptides. Red circles indicate a non-PfCSP reactive control TCR. (**d**) Percent *Pf* NF54 Th2R/T* aa 311-325, aa 315-329, and aa 319-333 peptide-reactive TCRs in each donor. Brackets link donors with TCRs restricted to the same HLA-DR allele. (**e**) Percent Dd2 cross-reactive TCRs among NF54 PfCSP aa 319-333 binders. Absolute TCR counts are provided above the bars. (**f**) *TRBV* segment usage of cross-reactive and non-cross-reactive aa 319-333 binders. (**g**) *TRA* and *TRB* V and J segment and CDR3 sequences among members of TCR similarity cluster 656. Amino acid mismatches in CDR3s of non-cross-reactive compared to the Dd2 cross-reactive TCR are highlighted in orange. (**c**) Results from one representative out of three independent experiments are shown.

## DISCUSSION

Despite their importance for humoral immune responses, little is known about the T_FH_ cell response to *Pf* and the sporozoite vaccine target PfCSP. Our single-cell based TCR repertoire analysis platform and algorithm to identify convergent TCRs provides unique first insights in the clonal dynamics, phenotype, TCR features, HLA restriction and epitope fine specificity of the human anti-sporozoite cT_FH_ cell response. Without antigen-based enrichment, we detected PfCSP-reactive T cell clones at high frequency in the activated cT_FH_ repertoire upon PfSPZ Vaccine immunization, demonstrating the immunodominant role of PfCSP among the plethora of *Pf* antigens. Successive immunizations were required to induce detectable PfCSP-specific cT_FH_ cell responses compared to the large number of non-PfCSP specific cT_FH_ cell clones that dominated in the early immunization phase. The paucity of cT_FH_ cell clones that were detected at multiple time points indicates that the successive immunizations diversified the repertoire through the recruitment of new cT_FH_ cell clones, comparable to the dynamic turnover of activated cT_FH_ cells after repeated influenza vaccinations^29^. Similar to natural *Pf* infections, the cT_FH_ cell response was skewed towards a T_H_1 phenotype^30^. Although the dominance of cT_FH_1 cells over cT_FH_2 cells has been shown to negatively impact humoral immunity, it remains to be determined whether this correlates with reduced protection or whether cT_FH_1 contribute to protection through the production of IFNγ^30–33^. The detection of numerous PfCSP-reactive cT_FH_ cell clones with mixed cT_FH_1 and cT_FH_2 phenotypes suggests that both effector functions co-exist among cells with the same antigen specificity in response to PfSPZ Vaccine immunization. Adjuvants may be used to shift the cT_FH_1 to cT_FH_2 ratio in vaccine formulations with recombinant PfCSP to define which cT_FH_ phenotype correlates with protective humoral immunity and vaccine efficacy.

Historic data, based on bulk CD4^+^ T cell analyses including limiting dilution experiments, have established Th2R/T* as immunodominant region containing epitopes that are universally presented on diverse HLA-DR molecules^21–23,34,35^ in contrast to T1 with reported presentation on a limited number of HLA molecules, including DRB1*04:01, DRB1*11:01 and DQB1*06:03 and DQA1*01:02^24,35,36^. Our data demonstrate the immunodominance of aa 311-333 in the Th2R/T* region over T1 at single-cell level for cT_FH_ cells and identify an epitope in the C-linker region that induced weak cT_FH_ responses in the context of rare HLA-DQ MHC complexes. Likely due to its reported lower immunogenicity, cT_FH_ cells with specificity for CS.T3, a known C-CSP epitope near the GPI anchor, were not detected in any of the donors^37,38^. Thus, efficient presentation on a wide variety of HLA-DR molecules compared to rare HLA-DQ molecules explains the abundance of anti-Th2R/T* cT_FH_ cells over any other epitope specificity.

Presumably due to absence of any of the reported T1-presenting alleles in all other donors except D3, which expressed HLA DQB1*06:03 and DQA1*01:02, anti-T1 cT_FH_ cells were detected only in D3^35,36^. T1-specific TCRs showed low CDR3 sequence similarity but enrichment of *TRBV30* suggesting that this segment plays an important role in TCR binding to the specific T1-MHC complex via CDR1 or CDR2. However, *TRBV30* abundance does not seem to influence the anti-T1 cT_FH_ response, since D1, D2 and D5 with the same *TRBV30* allele and similar *TRBV30* usage frequency before immunization compared to D3 lacked T1-specific TCRs. Whether the overall low generation probability of T1-specific TCRs contributes to the paucity of T1-reactive cT_FH_ cells remains unclear, but detection of equal numbers of anti-T1 and anti-Th2R/T* TCRs in D3 suggests that TCR repertoire diversity or antigen processing does not hinder the generation of anti-T1 cT_FH_ cells responses in donors with the presenting HLA complex.

Although T1 in the N-terminal junction is highly conserved and overlaps with epitopes of the most potent human anti-PfCSP antibodies that have been reported so far^11,12,14,39^, the inefficient targeting of T1 due to HLA restrictions limits its potential as T helper cell epitope in vaccine design^36^. In contrast, the universal HLA-DR presentation properties of Th2R/T* peptides and high generation probability of anti-Th2R/T* TCRs that are frequently encoded by *TRBV20-1* define this region as “supertope” that elicits strong convergent CD4^+^ T cell responses. Bulk cell analyses have shown that strong anti-Th2R/T* responses are also seen in non-European individuals^9,21,40^. Nevertheless, future studies at single-cell level should confirm that the potency of the aa 311-333 peptide as T helper cell epitope is not limited by genotype differences.

PfCSP-subunit vaccine designs may benefit from including only aa 311-333 within the Th2R/T* region rather than the complete C-terminus to avoid the induction of strong antibody responses against non-protective conformational epitopes in the aTSR domain^41^. Nevertheless, our data show that the anti-Th2R/T* T helper cell response will be limited by strain specificity. Cross-reactivity was only observed for TCRs against a more conserved sub-epitope covering aa 319-333. TCRs with this specificity were not identified in all individuals (presumably due to MHC-II genotype restrictions) and cross-reactivity was limited to a single strain with little sequence divergence compared to the PfSPZ Vaccine strain NF54. The observation that even highly similar TCRs differed in their ability to cross-react with the Dd2 and NF54-derived aa 319-333 peptides that vary by only one aa illustrates the sensitivity of the TCRs to subtle aa differences in their target epitopes. Due to the high degree of sequence polymorphisms in natural parasite populations, boosting of the T_FH_ cell response by non-vaccine strains will be inefficient and limited to individuals with potent responses against the aa 319-333 epitope. To what degree natural boosting increases vaccine efficacy or whether the strength of the anti-aa 311-333 T_FH_ cell response will be sufficient to promote long-term protection remains to be determined.

Our study provides fundamental insights in the clonal evolution of cT_FH_ cell response to *Pf* and highlights the power and importance of functional TCR repertoire studies at monoclonal level. The data define the potential of T cell target epitopes and identify factors that may constrain vaccine responses based on limitations in TCR repertoire diversity, HLA-restrictions, or cell phenotype. The relevance of our findings extends to anti-PfCSP CD4^+^ T cell responses beyond the cT_FH_ cell subset and may contribute to the design of an improved PfCSP-based malaria vaccine.

## METHODS

### Human immunization trial with PfSPZ Vaccine

The MAVACHE A clinical trial is registered in the EudraCT database under 2015-005123-11 and under the ClinicalTrials.gov number NCT02704533. Ethical approval was granted by the ethics committee of the medical faculty and the university clinics of the University of Tübingen. The study with healthy European volunteers with no history of malaria or HIV was carried out according to the principles of the Declaration of Helsinki. After signed informed consent, peripheral blood samples were collected three days before the first immunization and seven days after each of three immunizations with 9 × 105 radiation-attenuated, aseptic, purified, cryopreserved *Pf* NF54 sporozoites (Sanaria® PfSPZ Vaccine) on days 1, 8, and 29. PBMCs were isolated using Ficoll density gradient centrifugation, frozen and stored in liquid nitrogen until further use.

### Anti-PfCSP antibody ELISA

ELISAs were performed as described previously^42^. In brief, high-binding 96-well ELISA plates were coated with NF54 PfCSP (EMBL Protein Expression and Purification Core Facility, Heidelberg, Germany; 0.4 μg/ml) overnight at 4 °C. Plates were washed three times with deionized water, blocked with 4% BSA and 0.01% Tween in PBS for 1 h at room temperature, and washed again. Plasma samples diluted in PBS containing 1% BSA and 0.05% Tween were added for 1.5 h at room temperature. Plates were washed, incubated for 1 h with horseradish peroxidase conjugated goat anti-human IgG or IgA antibody diluted in PBS with 1% BSA and 0.05% Tween, washed again and incubated with ABTS solution (Roche). The reaction was stopped with 1 μl/ml H_2_O_2_. Absorbance at OD_405_ was measured with a M1000 Pro plate reader (Tecan).

### Flow cytometry and single-cell sorting

For flow cytometric cell analyses and cell sorting, freshly thawed and washed PBMCs (5×106) were incubated for 30-60 min at 4 °C in the dark with 100 μl staining cocktail, containing the following fluorescently-labelled antibodies diluted in FACS buffer (2% FBS in PBS): CD4-APC-Cy7 (clone A161A1; BioLegend), CD8a-Alexa700 (clone SK1; BioLegend); CD45RA-BV510 (clone HI100; BioLegend); PD-1-BV605 (clone EH12.2H7; BioLegend), ICOS-PE-Cy7 (clone C398.4A; BioLegend), CXCR5-Alexa647 (clone RF8B2; BD Biosciences), CXCR3-BV421 (clone G025H7; BioLegend), CCR6-PE (clone G034E3; BioLegend) and CCR7-BV710 (clone G043H7; BioLegend). Subsequently, cells were washed with FACS buffer and incubated with the live-dead marker 7-aminoactinomycin D (7-AAD; Thermo Fisher Scientific) for 10 min at 4 °C in the dark. Flow cytometric analyses and indexed single-cell sorting were performed using a FACS AriaIII (BD Biosciences) and FACSDiva software (version 8.0.1) as described^19^. In brief, single activated circulating 7-AAD-CD3+ CD4+ CD45RA-CXCR5+ PD-1hi ICOS+ cT_FH_ cells were sorted into 384-well plates containing 2 μl random hexamer primer (RHP) mix (consisting of nuclease free water, PBS, RNAsin, NP-40, DTT, random hexamer primers), immediately sealed, frozen on dry ice, and stored at −80 °C until further processing.

### cDNA synthesis and TCR gene amplification

cDNA synthesis and TCR gene amplification was performed as described previously^19^. In brief, cDNA was prepared in a total volume of 4 μl in the original sort plates using random hexamer primers. TRA and TRB transcripts were amplified in separate semi-nested PCRs in a total volume of 10 μl using pre-diluted cDNA as template. TRA and TRB amplification used TRAV and TRBV specific primer sets and respective constant region primers. Two independent primary PCR reactions were performed to increase the amplification efficiency^43^. Secondary PCRs were set up using 0.5 μl of each of the two independent first PCRs as template. A common linker sequence^44^ attached to all primary PCR TRAV and TRBV primers served as binding site for a universal forward primer in the secondary PCR enabling amplification in combination with nested constant region primers. Forward and reverse primers used in the secondary PCR contained 16 bp-long row- and column-specific barcodes to preserve TRA and TRB pairing information at single-cell level upon pooling of all amplicons for next-generation sequencing using Illumina MiSeq 2×300 (Eurofins Genomics)^45^. All pipetting steps were carried out on a Tecan EVO200 automation platform. cDNA synthesis and PCR reactions were performed using Eppendorf MasterCyclers.

### TCR sequence analysis

TCR sequence assembly and segment annotation was performed by a modified version of the automated immunoglobulin (Ig) gene analysis pipeline sciReptor^46^. In brief, sciReptor used PANDAseq^47^ to assemble paired end Illumina sequencing reads with a minimum overlap of 50 bp into assembled reads with a maximum and minimum length of 550 and 300 bp, respectively, and a minimum quality score of 0.8. Reads were assigned to corresponding wells based on their positional barcodes. V, D and J segments were annotated using IgBLAST^48^ and the Ensembl Genome Browser^49^, and CDR3 sequence information were obtained. Flow cytometric index data and TCR sequence information were linked and stored in a MariaDB database. Downstream sequence analyses and data visualization were performed using custom R scripts. T cell clones were defined as cells having the same TRBV and TRBJ segment as well as identical TRB CDR3 nucleotide sequence. Repertoire diversity was characterized by normalized Shannon Wiener index^50^ estimated by bootstrap with 10,000 replicates. Repertoire overlap was calculated as an average fraction of shared clonotypes between subsampled repertoires (N=50) calculated in 10,000 repetitions. TCR generation probabilities were calculated as a product of TRA and TRB generation probabilities estimated using OLGA^51^.

### TCR clustering

To identify clusters of convergent TCRs that are likely to recognize the same peptide, a sequence similarity analysis framework was developed (Supplementary Figure 3). The clustering is based on pairwise comparisons of all TCRs by aligning their CDR1-3 and calculating the alignment scores using BLOSUM62 substitution matrix reflecting evolutionary amino acid interchangeability. The three alignment scores (CDR1, CDR2 and CDR3) are logistically transformed into a single score (BL-score, BLOSUM-logistic) ranging from 0 to 1, where 1 corresponds to the highest probability of specificity match and 0 to the lowest. Because the CDRs differ in their impact on TCR specificity, they are weighted differently in the logistic function. The weights were established using available data sets of TCRs with known specificity (VDJdb^52^ and IEDB^53^). An optimal cut-off BL-score for clustering was also inferred from the training data set. The clustering was validated using data (20%) not included in the training set and compared to available tools^54–56^ to assess its performance (Supplementary Figure 4). The code is available at https://github.com/obrzts/BLscore.

### TCR expression cloning

The cloning and stable expression of TCRs in TCR^neg^ CD3^+^ CD4^+^ Jurkat76 cells (J76-CD4) was performed as described previously^19^. In brief, full-length *TRA* and *TRB* genes were cloned into retroviral pMSCV-PImC expression vectors. Retroviral particles were harvested from the supernatants of Phoenix Ampho cells (ATCC) cultured in DMEM GlutaMAX medium (Life Technologies) with 10% heat-inactivated FBS at 37 °C and 5% CO_2_ after transfection with the TCR-expression vectors using 2.5 M CaCl_2_ and used for the transduction of J76-CD4 T cells. TCR (clone IP26; BioLegend) expression on live cells (LIVE/DEAD Fixabable Near-IR Dead Cell Stain; Invitrogen) was confirmed by flow cytometry after 7 days of 0.8 μg/ml puromycin dihydrochloride selection.

### TCR reactivity tests

TCR reactivity tests were performed as described^19^. In brief, 2.13×10^5^ Epstein-Barr-Virus (EBV) immortalized B cells were seeded in 96-well tissue culture plates (U-bottom) in AIM-V medium and loaded with 2.5 μg/ml peptide for 18 h at 37 °C and 5% CO_2_ before addition of 4.27×10^5^ TCR-transgenic Jurkat76 cells. For HLA-blocking experiments, 10 μg/ml HLA-DR (clone L243, BioLegend), HLA-DQ (clone SPVL3, Abeomics) or HLA-DP (clone B7/21, Leinco Technologies) blocking antibody were added together with the T cells^57^. Supernatants were harvested 24 hours later and IL-2 concentrations were determined using serially diluted supernatants and the human IL-2 ELISA MAX Deluxe kit (BioLegend) adapted to 384-well plate format using one quarter of the recommended reaction volumes. Optical density (OD) values were measured at 450 nm with M1000 Pro plate reader (Tecan).

### Peptide-MHC binding prediction

HLA-typing was performed by DKMS Life Science Lab GmbH. MHC-II sharing between donors was determined by the presence of MHC-II alpha and beta chain pairs (DQ, DP) or beta chains only (DR) with identical aa sequence in the peptide binding groove (α1 and β1 domain). For in silico analyses, NetMHCIIpan-4.0^25^, Sturniolo^26^, and MixMHC2pred^27^ were used with default parameters; strong and weak binding was defined by a score within the top 2% and 10% percentile, respectively, of the binding scores computed for random peptides and a given MHC-II allele. MHCnuggets^28^ was used with default parameters and strong and weak binding was defined by a predicted IC50 below 50 and 500 nmol/L, respectively.

### Statistical analyses

Statistics were calculated using GraphPad Prism (Version 8.1.2) and R (Version 4.0.4). The corresponding statistical test used for each experiment is stated in the figure legend (*P < 0.05; **P < 0.01; ***P < 0.001; ****P < 0.0001).

## DATA AVAILABILITY STATEMENT

The datasets generated during and/or analyzed during the current study are available on Zenodo, https://doi.org/10.5281/zenodo.4542509.

## ACKNOWLEDGEMENTS

The authors thank Dorien Foster, Julia Gärtner and Claudia Winter (DKFZ) for experimental assistance, Thomas Höfer, Matthias Günther, Christian E. Busse and Francisco Arcila Salamanca (DKFZ) for bioinformatics support. We thank the team of the EMBL Protein Expression and Purification Core Facility, especially Kim Remans, for their support. We further thank the vaccine trial participants and all members of the clinical trial platform for their contribution and commitment to vaccine research. The clinical trial was funded by the Deutsches Zentrum für Infektionsforschung (German Center for Infection Research – DZIF), award number TTU 03.902 to B.M. Manufacture of PfSPZ Vaccine was funded in part by the National Institute of Allergy and Infectious Diseases of the National Institutes of Health under SBIR award numbers 5R44AI058375 and 5R44AI055229.

## AUTHORS CONTRIBUTION

I.W. and H.W. designed experiments. I.W., A.O., J.P., R.H. performed experiments. I.W., A.O. and H.W. analyzed experimental data. I.W. and A.O. performed statistical analyses. A.O. developed bioinformatics tools. S.C., B.K.L.S., S.L.H., P.G.K. and B.M. produced the vaccine and/or provided clinical trial samples. B.M. and H.W. conceived the study. I.W., A.O. and H.W. wrote the manuscript.

