## Supplemental Figures 1-5 and Supplemental Tables 2-4 for "Clonal evolution and specificity of the human T follicular helper cell response to *Plasmodium falciparum* circumsporozoite protein"

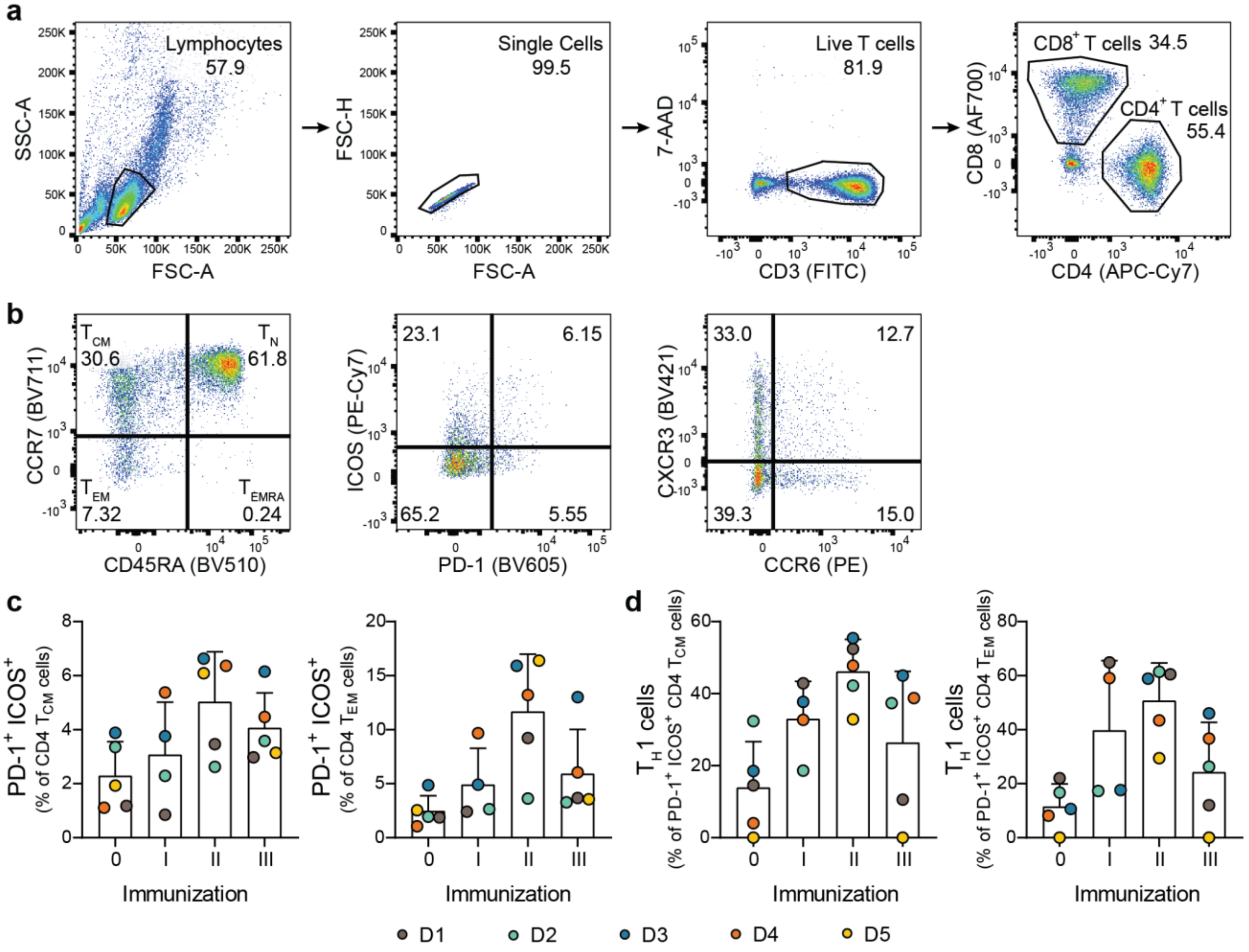


**Supplementary Figure 1: Flow cytometric analysis of $CD4^+$ T cell responses to repeated PfSPZ Vaccine immunizations.** (**a**) Representative gating strategy for the identification of $CD3^+CD4^+$ T cells. Dead cells were excluded by 7-AAD stain. (**b**) Representative gating strategy for the analysis of $CD4^+$ $T_{CM}$ ($CD45RA^-$ $CCR7^+$) and $CD4^+$ $T_{EM}$ ($CD45RA^-CCR7^-$) cells. (**c,d**) Quantification of activated ($PD\text{-}1^+ICOS^+$) (**c**) or $T_H1$ ($CXCR3^+CCR6^-$) $CD4^+$ $T_{CM}$ and $T_{EM}$ cells. Dots represent individual donors, bars show mean and SD.

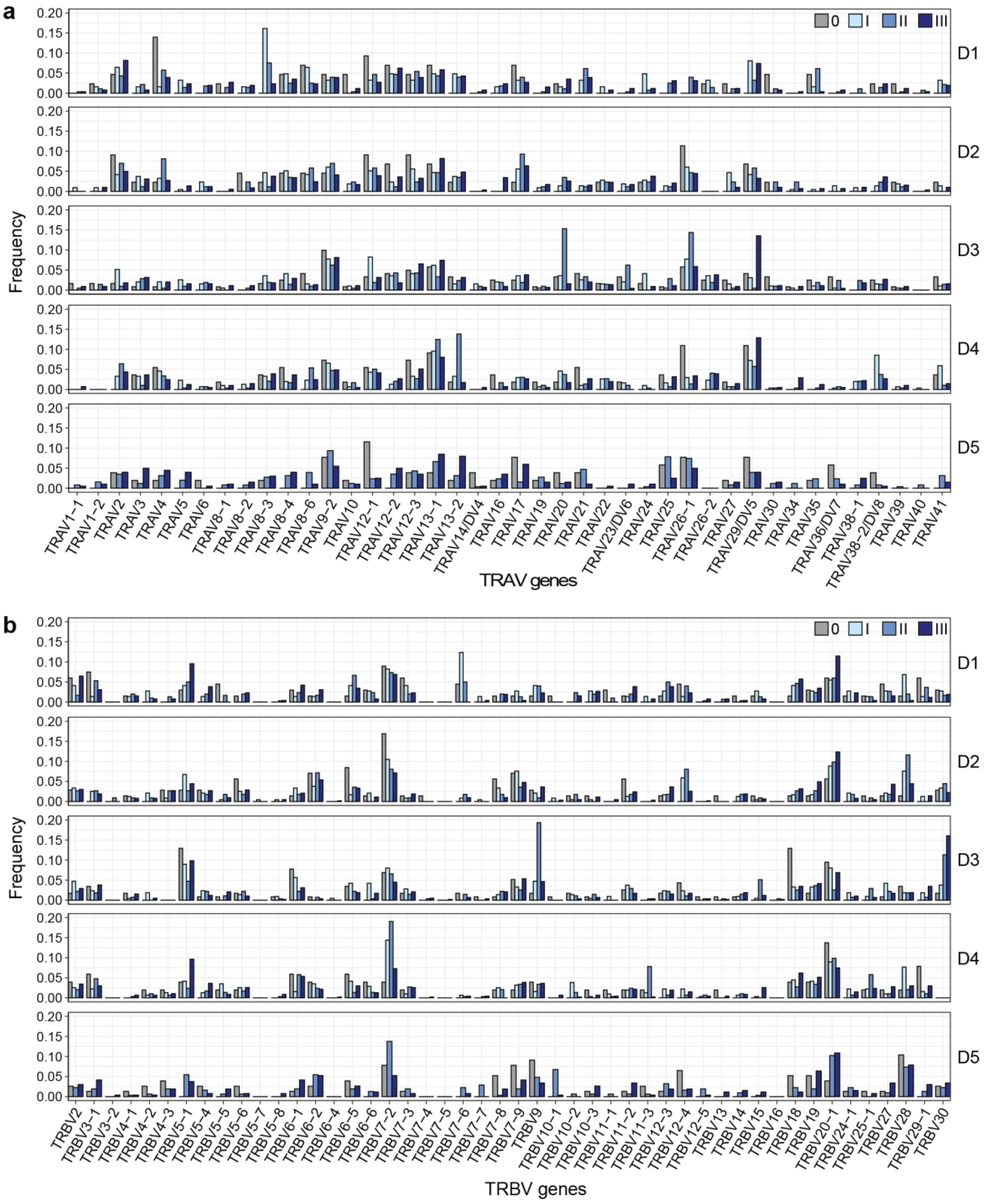


**Supplementary Figure 2: Changes in TR V segment usage after PfSPZ Vaccine immunization.** Quantification of *TRAV* (**a**) and *TRBV* (**b**) segment usage before (0) and after PfSPZ Vaccine immunizations I-III.

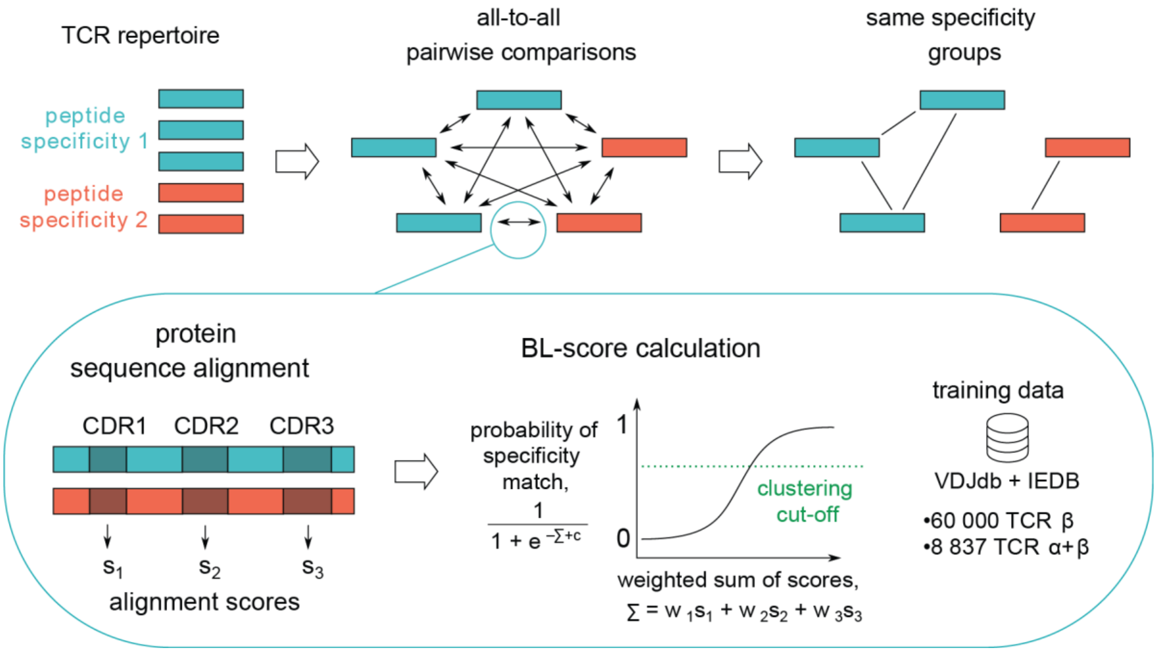


**Supplementary Figure 3: Scheme for clustering of similar TCRs with the same peptide specificity.** All TCRs in a given repertoire are compared pairwise. First, CDR1-3 of each TCR pair are aligned and the alignment scores are calculated using BLOSUM62 matrix. Next, probability of specificity match (BL-score) is calculated as a logistic function of weighted sum of the three alignment scores. The weights and the BL-score cut-off for the clustering were learned from the TCR data[52,53] with known peptide specificity. The clusters are defined by applying the cut-off and detecting the groups of connected TCRs.

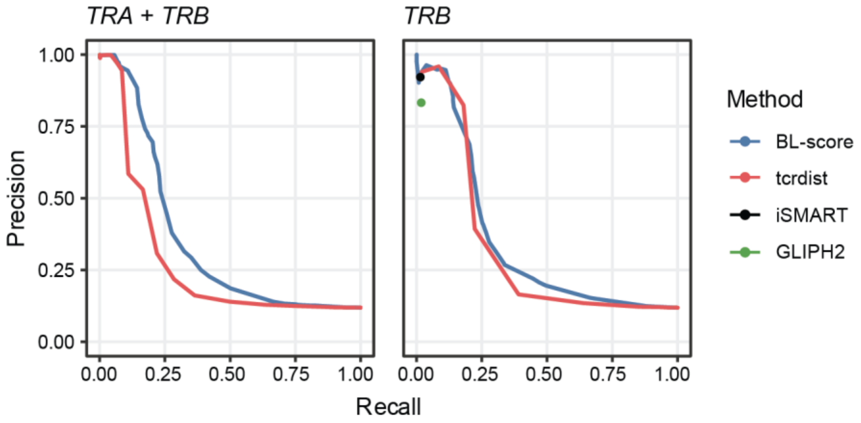


**Supplementary Figure 4: Performance of TCR clustering methods applied to *TRA* + *TRB* or *TRB* datasets.** Clustering using BL-score and tcrdist[54] was performed using varying cut-offs; iSMART[55] and GLIPH2[56] were used with the default parameters. Precision was calculated as a fraction of TCR pairs with the same specificity among all pairs of TCRs that were clustered together. Recall was calculated as a fraction of clustered TCR pairs among all pairs of TCRs with the same specificity. The comparison was done using 20% of the data not included in BL-score clustering training set.

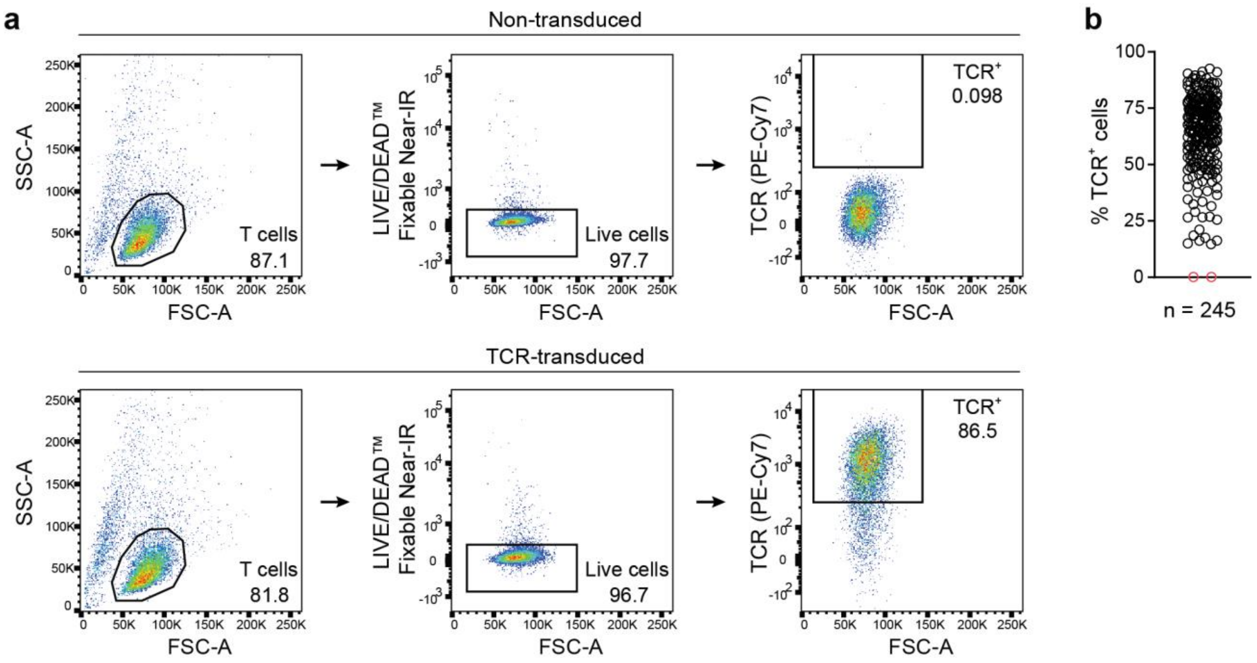


**Supplementary Figure 5: TCR expression on Jurkat76 T cells upon TCR-transduction.** Flow cytometric analysis of Jurkat76 T cells before and after TCR-transduction. Representative gating strategy for the identification of live TCR+ T cells (**a**) and quantification for all 245 analyzed TCRs (**b**). Each dot indicates a different TCR, the non-transduced control is shown in red.

SUPPLEMENTARY TABLES

**Supplementary Table 1: TCR sequence features and reactivity data.**

**Supplementary Table 2: List of PfCSP peptides used for T cell stimulation.**

| | Sequence | Position | Name |
|---|---|---|---|
| N-CSP peptide pool | MMRKLAILSVSSFLF | 1-15 | CSP1-15 |
| | LAILSVSSFLFVEAL | 5-19 | CSP5-19 |
| | SVSSFLFVEALFQEY | 9-23 | CSP9-23 |
| | FLFVEALFQEYQCYG | 13-27 | CSP13-27 |
| | EALFQEYQCYGSSSN | 17-31 | CSP17-31 |
| | QEYQCYGSSSNTRVL | 21-35 | CSP21-35 |
| | CYGSSSNTRVLNELN | 25-39 | CSP25-39 |
| | SSNTRVLNELNYDNA | 29-43 | CSP29-43 |
| | RVLNELNYDNAGTNL | 33-47 | CSP33-47 |
| | ELNYDNAGTNLYNEL | 37-51 | CSP37-51 |
| | DNAGTNLYNELEMNY | 41-55 | CSP41-55 |
| | TNLYNELEMNYYGKQ | 45-59 | CSP45-59 |
| | NELEMNYYGKQENWY | 49-63 | CSP49-63 |
| | MNYYGKQENWYSLKK | 53-67 | CSP53-67 |
| | GKQENWYSLKKNSRS | 57-71 | CSP57-71 |
| | ENWYSLKKNSRSLGE | 61-75 | CSP61-75 |
| | SLKKNSRSLGENDDG | 65-79 | CSP65-79 |
| | NSRSLGENDDGNNED | 69-83 | CSP69-83 |
| | LGENDDGNNEDNEKL | 73-87 | CSP73-87 |
| | DDGNNEDNEKLRKPK | 77-91 | CSP77-91 |
| | NEDNEKLRKPKHKKL | 81-95 | CSP81-95 |
| | EKLRKPKHKKLKQPA | 85-99 | CSP85-99 |
| | KPKHKKLKQPADGNP | 89-103 | CSP89-103 |
| | KKLKQPADGNPDPNA | 93-107 | CSP93-107 |
| | QPADGNPDPNANPNV | 97-111 | CSP97-111 |
| | GNPDPNANPNVDPNA | 101-115 | CSP101-115 |
| | PNANPNVDPNANPNV | 105-119 | CSP105-119 |
| | PNVDPNANPNVDPNA | 109-123 | CSP109-123 |
| | NANPNANPNANPNAN | 257-271 | CSP257-271 |
| C-CSP peptide pool | NPNANPNANPNKNNQ | 263-277 | CSP263-277 |
| | NPNANPNKNNQGNGQ | 267-281 | CSP267-281 |
| | NPNKNNQGNGQGHNM | 271-285 | CSP271-285 |
| | NNQGNGQGHNMPNDP | 275-289 | CSP275-289 |
| | NGQGHNMPNDPNRNV | 279-293 | CSP279-293 |
| | HNMPNDPNRNVDENA | 283-297 | CSP283-297 |
| | NDPNRNVDENANANS | 287-301 | CSP287-301 |
| | RNVDENANANSAVKN | 291-305 | CSP291-305 |
| | ENANANSAVKNNNNE | 295-309 | CSP295-309 |
| | ANSAVKNNNNEEPSD | 299-313 | CSP299-313 |
| | VKNNNNEEPSDKHIK | 303-317 | CSP303-317 |
| | NNEEPSDKHIKEYLN | 307-321 | CSP307-321 |
| | PSDKHIKEYLNKIQN | 311-325 | CSP311-325 |
| | HIKEYLNKIQNSLST | 315-329 | CSP315-329 |
| | YLNKIQNSLSTEWSP | 319-333 | CSP319-333 |
| | IQNSLSTEWSPCSVT | 323-337 | CSP323-337 |

| | | | |
|---|---|---|---|
| | LSTEWSPCSVTCGNG | 327-341 | CSP327-341 |
| | WSPCSVTCGNGIQVR | 331-345 | CSP331-345 |
| | SVTCGNGIQVRIKPG | 335-349 | CSP335-349 |
| | GNGIQVRIKPGSANK | 339-353 | CSP339-353 |
| | QVRIKPGSANKPKDE | 343-357 | CSP343-357 |
| | KPGSANKPKDELDYA | 347-361 | CSP347-361 |
| | ANKPKDELDYANDIE | 351-365 | CSP351-365 |
| | KDELDYANDIEKKIC | 355-369 | CSP355-369 |
| | DYANDIEKKICKMEK | 359-373 | CSP359-373 |
| | DIEKKICKMEKCSSV | 363-377 | CSP363-377 |
| | KICKMEKCSSVFNVV | 367-381 | CSP367-381 |
| | MEKCSSVFNVVNSSI | 371-385 | CSP371-385 |
| | SSVFNVVNSSIGLIM | 375-389 | CSP375-389 |
| | SSIGLIMVLSFLFLN | 383-397 | CSP383-397 |
| Cross-reactivity | PSDKHIEQYLKKIQNSLSTEWSP | 311-333 | NF54_CSP311-333 |
| | PSDKHIEQYLKKIQYSLSTEWSP | 311-333 | 7G8_CSP311-333 |
| | PSDKHIEQYLKKIKNSISTEWSP | 311-333 | K1_CSP311-333 |
| | PSDKHIKEYLNKIQNSLSTEWSP | 311-333 | Dd2_CSP311-333 |

**Supplementary Table 3: Prediction of pMHC binding for PfCSP peptides.**

| Donor | Epitope | Locus | MHC molecule | Experimental | netMHCIIpan | MHCnuggets | Sturniolo | mixMHCIIpred |
|---|---|---|---|---|---|---|---|---|
| D1 | Th2R | DR | DRB1*07:01P | potentially | no | weak | no | weak |
| D1 | Th2R | DR | DRB1*14:01P | | no | weak | - | - |
| D1 | Th2R | DR | DRB3*02:02P | | no | strong | - | weak |
| D1 | Th2R | DR | DRB4*01:01P | | weak | strong | - | strong |
| D1 | Th2R | DQ | DQA1*01:01P-DQB1*02:01P | | weak | no | - | - |
| D1 | Th2R | DQ | DQA1*01:01P-DQB1*05:03P | | weak | no | - | - |
| D1 | Th2R | DQ | DQA1*02:01P-DQB1*02:01P | potentially | no | no | - | weak |
| D1 | Th2R | DQ | DQA1*02:01P-DQB1*05:03P | | weak | weak | - | - |
| D2 | Th2R | DR | DRB1*04:02P | potentially | strong | weak | no | - |
| D2 | Th2R | DR | DRB1*07:01P | | no | weak | no | weak |
| D2 | Th2R | DR | DRB4*01:01P | | weak | strong | - | strong |
| D2 | Th2R | DP | DPA1*02:01P-DPB1*01:01P | potentially | weak | no | - | - |
| D2 | Th2R | DP | DPA1*02:01P-DPB1*14:01P | | no | weak | - | - |
| D3 | Th2R | DR | DRB1*07:01P | | no | weak | no | weak |
| D3 | Th2R | DR | DRB1*15:01P | potentially | strong | weak | no | strong |
| D3 | Th2R | DR | DRB4*01:01P | | weak | strong | - | strong |
| D3 | Th2R | DR | DRB5*01:01P | potentially | no | weak | no | weak |
| D4 | Th2R | DR | DRB1*01:01P | | no | strong | no | weak |
| D4 | Th2R | DR | DRB1*15:01P | potentially | strong | weak | no | strong |
| D4 | Th2R | DR | DRB5*01:01P | potentially | no | weak | no | weak |
| D5 | Th2R | DR | DRB1*04:02P | potentially | strong | weak | no | - |
| D5 | Th2R | DR | DRB1*11:04P | | weak | weak | no | weak |
| D5 | Th2R | DR | DRB4*01:01P | | weak | strong | - | strong |
| D5 | Th2R | DR | DRB3*02:02P | | no | strong | - | weak |
| D3 | T1 | DQ | DQA1*01:02P-DQB1*02:01P | potentially | weak | no | - | - |
| D3 | T1 | DQ | DQA1*01:02P-DQB1*06:03P | potentially | weak | no | - | - |
| D3 | T1 | DQ | DQA1*02:01P-DQB1*02:01P | | no | no | - | no |
| D3 | T1 | DQ | DQA1*02:01P-DQB1*06:03P | potentially | no | no | - | - |
| D5 | C-linker | DQ | DQA1*03:01P-DQB1*03:01P | potentially | weak | weak | - | strong |
| D5 | C-linker | DQ | DQA1*03:01P-DQB1*03:02P | | weak | no | - | - |
| D5 | C-linker | DQ | DQA1*05:01P-DQB1*03:01P | potentially | weak | weak | - | no |
| D5 | C-linker | DQ | DQA1*05:01P-DQB1*03:02P | potentially | no | no | - | - |

**Supplementary Table 4: TCR sequence features of Th2R/T*-specific TCRs grouped by TCR similarity.** Residues conserved within a cluster are shown in bold.

| | | TCR beta chain | | | TCR alpha chain | | |
|---|---|---|---|---|---|---|---|
| cluster ID | Donor | V | J | CDR3 | V | J | CDR3 |
| 1137 | D1 | 5-1 | 2-3 | **ASS**LVE**GPETQY** | 5 | 15 | **A**CRT**QAGTALI** |
| | D1 | 5-1 | 2-5 | **ASS**HIQ**GPETQY** | 5 | 15 | **A**EKN**QAGTALI** |
| 580 | D2 | 20-1 | 1-5 | **SA**SPP**R**S**N**QPQH | 24 | 48 | **AF**T**NFGNEKLT** |
| | D2 | 20-1 | 1-6 | **SA**PGQ**R**G**N**SPLH | 24 | 48 | **AF**N**NFGNEKLT** |
| | D2 | 20-1 | 1-6 | **SA**PTG**R**V**N**SPLH | 24 | 48 | **AF**S**NFGNEKLT** |
| | D2 | 20-1 | 1-1 | **SA**PRG**R**M**N**TEAF | 24 | 48 | **AF**T**NFGNEKLT** |
| 656 | D2 | 20-1 | 1-2 | **SA**PRR**R**A**NYGYT** | 24 | 48 | **A**WG**NFGNEKLT** |
| | D2 | 20-1 | 1-2 | **SA**TPR**R**V**NYGYT** | 24 | 48 | **A**FS**NFGNEKLT** |
| | D2 | 20-1 | 1-2 | **SA**TPH**R**A**NYGYT** | 24 | 48 | **A**RT**NFGNEKLT** |
| | D2 | 20-1 | 1-2 | **SA**TGR**R**A**NYGYT** | 24 | 48 | **A**ST**NFGNEKLT** |
| 286 | D5 | 20-1 | 2-7 | **SARD**S**GRSSYEQY** | 16 | 31 | **ALR**GAR**L**M |
| | D2 | 20-1 | 2-7 | **SARD**G**GRSSYEQY** | 16 | 23 | **ALR**GQGGK**L**I |
| | D5 | 20-1 | 2-7 | **SARD**G**GRSSYEQY** | 16 | 58 | **ALR**VGGSR**L**T |
| | D2 | 20-1 | 2-7 | **SARD**G**GRSSYEQY** | 16 | 31 | **ALR**GSAR**L**M |
| 1172 | D3 | 5-1 | 2-7 | **AS**TP**GGR**AEE**QY** | 29/DV5 | 48 | **AAR**S**NFGNEKLT** |
| | D3 | 5-1 | 2-3 | **AS**SP**GGR**SDT**QY** | 29/DV5 | 48 | **AAR**T**NFGNEKLT** |
| | D4 | 5-1 | 2-3 | **AS**SS**GGR**SDT**QY** | 29/DV5 | 48 | **AAR**T**NFGNEKLT** |
| 1745 | D3 | 7-8 | 1-6 | **ASS**PH**RAGDSPLH** | 13-2 | 28 | **AENRR**A**GSY**QL**T** |
| | D3 | 7-8 | 1-6 | **ASS**RE**RAGDSPLH** | 13-2 | 6 | **AENRR**G**GSY**IP**T** |
| 1169 | D4 | 7-9 | 2-5 | **ASSPGGR**KE**TQY** | 29/DV5 | 45 | **A**AHT**DSGGGADGLT** |
| | D3 | 7-9 | 2-3 | **ASSPGGR**AD**TQY** | 25 | 45 | **A**GNR**DSGGGADGLT** |
| 1652 | D4 | 9 | 1-2 | **ASSVAGEDYGYT** | 12-2 | 22 | **AV**I**L**S**GSARQLT** |
| | D3 | 9 | 1-2 | **ASSVAGEDYGYT** | 12-2 | 22 | **AV**L**L**G**GSARQLT** |
| 1239 | D4 | 18 | 2-3 | **ASS**G**GGGRGTDTQY** | 29/DV5 | 43 | **AASARDMR** |
| | D3 | 18 | 2-3 | **ASS**P**GGGRGTDTQY** | 29/DV5 | 43 | **AASARDMR** |
